# Enhancing CRISPR Education: A Plug-And-Play Framework for Teaching Gene Editing in Plant Biology

**DOI:** 10.1101/2025.10.10.681630

**Authors:** Divya Jain, Faizan Ali, Gift Obunkukwu, Hyndavi Yammanuru, Jing Zou, Joshua Obeng, Kyla Danae Hughes, Laxmi Joshi, Madhavarapu Sudhakar, Oluwaseun Adeyemi, Peter Prestwich, Shahla Borzouei, Shivani Dharam, Shubh Pravat Singh Yadav, Mst Sumayae Khan, Rajni Parmar, Upama Adhikari, Ali Taheri, Robin Taylor, Vicki Caruana, Mary Williams, Sonali Roy

## Abstract

Modern molecular biology tools and technologies such as CRISPR have sped up scientific discovery. From an educational perspective, these advancements are both exciting and overwhelming. Educators shaping these future scientists face the ongoing challenge of staying compliant with the latest developments in molecular biology while finding effective ways to teach these discoveries. As the use of CRISPR gene editing technology continues to expand globally, there is an increasing need for a workforce that is both knowledgeable about its theoretical foundations and trained in its practical use. While advanced, technology-driven STEM courses have the potential to improve student retention, they are often lecture-heavy and lack intentional engagement strategies that support deeper learning. Moreover, agriculture is the second most impacted sector by this technology, yet there is a significant lack of teaching materials focused on CRISPR in plant biology. To address these gaps, we developed a framework for teaching gene editing that incorporates multiple engagement strategies beyond traditional lecture-based instruction. This framework was implemented over two semesters in an *Introduction to Gene Editing* course at Tennessee State University, offered to both undergraduate and graduate students enrolled in a degree in Agricultural Sciences. This manuscript outlines the various strategies used in the course including active learning, multimodal instructional approaches and experiential learning strategies that can be adopted in other classrooms to effectively teach gene editing. Survey-based results from the course indicate a measurable increase in student comfort with designing and executing CRISPR-Cas based experiments.

**Societal Impact Statement:** Plant biology lacks accessible teaching materials for CRISPR, a powerful gene-editing technology widely used to improve agriculture. We developed an engaging framework to teach CRISPR concepts to advanced undergraduates and graduate students in plant sciences, which can be readily adopted by other instructors. The approach increased self-reported confidence in students and comfort in explaining CRISPR. Since instructors often have limited time to design interactive lessons, this framework offers a ready-to-use, effective strategy that makes CRISPR more widely available in classrooms, ultimately strengthening CRISPR literacy in the future agricultural workforce.

## 1 INTRODUCTION

The area of plant biology has been tremendously impacted by gene-editing technologies, specifically CRISPR-Cas, which have made it possible to manipulate the genetics of crops and conduct ecological research more easily (Yuan *et al*., 2024). CRISPR, which stands for Clustered Regularly Interspaced Short Palindromic Repeats, functions in conjunction with CRISPR-associated (Cas) proteins to recognize and cleave specific DNA sequences, typically 20–30 nucleotides in length (Jinek *et al*., 2012). Originally discovered as part of the bacterial immune system, CRISPR sequences are located in unique regions of the bacterial genome, where they help detect and destroy foreign DNA, such as that from invading phages, before it can integrate into the host genome. Since its adaptation as a molecular tool, CRISPR-Cas has rapidly become widely used across the life sciences, enabling breakthroughs in medicine and agriculture, including the development of disease-resistant crops with enhanced yield and nutritional value (Ansori *et al*., 2023). As these technologies continue to evolve, there is an emerging need for plant biology education to stay on top of the advances. Conventional teaching methodologies that concentrate mainly on lectures and textbooks may not deliver the depth of knowledge and practical experience necessary for teaching CRISPR successfully. Some challenges for teaching CRISPR technology, include complex molecular mechanisms, specialized terminology, and abstract thinking that may not be fully covered in primary fields of study for biotechnology students from diverse backgrounds like animal science, plant breeding, and tissue culture.

This manuscript focuses on teaching techniques and advanced biotechnology pedagogies adopted into the Agricultural Science degree program at the Tennessee State University. The objective of the ‘Introduction to Gene Editing with CRISPR-Cas’ in Plant Biology course at Tennessee State University was to help students appreciate the broad impact of CRISPR technologies in addressing global agricultural challenges, using diverse tools and techniques to create a more engaging learning experience.

In addition to traditional lectures, this course was designed to incorporate active learning, multimodal teaching strategies, and experiential learning.

1. Active learning strategies in advanced STEM courses significantly enhance student engagement and retention over time (Alaagib *et al*., 2019; Sudarmika *et al*., 2020; Zeng *et al*., 2021). Research has also proven that active learning improves standardized test scores compared to traditional lectures alone (Freeman *et al*., 2014). From the student’s perspective, active learning leads to more curiosity-based learning because they feel motivated to ask questions, give their input and critique the experimental approaches. For instructors, critical response from students in real time helps them adjust their teaching methodologies consistently and present updated research materials. The interactive learning strategy can also help in reducing the number of students who drop out of STEM courses because they feel bored or find the topics too hard (Olson & Riordan, 2012).
2. Multimodal teaching strategies enhance student engagement and comprehension by incorporating diverse formats of learning. These strategies include: writing blog posts to encourage reflection and improve science communication skills, analyzing film and television media as case studies to critically evaluate public representations of science, and using online teaching platforms commonly used at universities to initiate structured debates that develop critical thinking and collaborative learning. Together, these approaches accommodate different learning styles and make complex scientific concepts more accessible and relevant.
3. Experiential learning, which stresses active engagement in scientific processes, is important for teaching CRISPR because it encourages critical thinking and helps students to build practical skills that are needed for success in plant biotechnology and genetic research(Wolyniak *et al*., 2019). These include hands on molecular biology experiments such as cloning, PCR and gel electrophoresis; the use of digital tools to design primers and guide RNAs; plant phenotyping techniques including growth on sterile media and in soil. Using plants as model organisms helps the teaching of CRISPR technologies. Compared to animal systems, plants typically do not require extensive regulatory approvals for genetic manipulation, have relatively short life cycles, and often exhibit visible phenotypic changes when edited which is easy for students to observe. For example, in the model flowering plant *Arabidopsis thaliana*, genes involved in pigment biosynthesis such as *PHYTOENE DESATURASE3* (*PDS*3), which encodes a key rate-limiting enzyme in carotenoid biosynthesis, can be targeted with CRISPR to produce an albino phenotype, providing an easily observable phenotype for students (Mills *et al*., 2021). In addition, using plants in CRISPR exercises helps students learn how gene editing can be used to address some of the world’s foremost issues, such as increasing the food supply, ameliorating the effects of climate change on crops and ecosystems, promoting food security on a global scale, and even growing food for space explorations (Hubbard, 2024).

## 2 MATERIALS AND METHODS

### 2.1. Pedagogical approaches

The interactive classroom activities and student evaluation of performance are described in the text of this article. The lab practical activities including DNA isolation, PCR, and plant growth and transformation are described in detail in the Supplemental materials.

### 2.2. Student assessment of learning: Student surveys

To understand student perceptions, engagement levels, and learning outcomes related to the innovative pedagogies used in this course, a mixed-methods approach was employed. Data collection included pre- and post-surveys and focus group discussions with students enrolled across two cohorts of the CRISPR course at Tennessee State University. All students participated in a focus group held during the last scheduled hour of the course. In addition, 20 of the 23 students complete pre-surveys, 18 of 23 completed post-surveys with a total of 15 students having matched data across the pre- and post-survey administration. All students were Agricultural Sciences majors.

All raw responses were exclusively accessible to a third-party evaluator, who removed identifying information from the data and provided only aggregated summary data to the course instructor and the research team. The surveys comprised a combination of quantitative and open-ended questions to investigate the influence of the course on students’ understanding, perceptions, and comfort levels related to genetic engineering, particularly CRISPR-Cas technology. A final aspect of the surveys was to evaluate changes in students’ comfort levels with various gene-editing techniques before and after the course. Post-surveys were also used to understand students’ perceived ability to explain gene-editing concepts to others, recommendation of the course, and if the course increased their interest in genetic engineering. Semi-structured focus group sessions to gather in-depth qualitative data on students’ experiences, perceptions, and learning outcomes were also conducted across two academic years.

## 3 RESULTS

The Introduction to Gene Editing using CRISPR-Cas course used a variety of teaching methods to deliver content including lecture-based teaching (**Figure 1**). The instructor presented 1-1.5 hours of lecture content using no more than 15-25 Microsoft Office PowerPoint slides that integrated material available on the open source CRISPRPedia as well as research articles available on Pubmed or Google Scholar. The slides included both essential content and material labeled ‘good to know’ - information that was interesting and relevant, but too advanced for the current course level. PowerPoint provided a visual aid while the instructor used lectures to explain the components of CRISPR technology and how the Cas9 complex creates gene edits. Lectures were intentionally aligned with practical sessions, with key molecular biology concepts introduced in advance of their application. For example, topics such as reading a plasmid map and the principles of Golden Gate cloning were taught prior to performing the cloning experiments. Additional slides were used as needed to explain related concepts such as the process of bacterial transformation.

**Figure 1.**
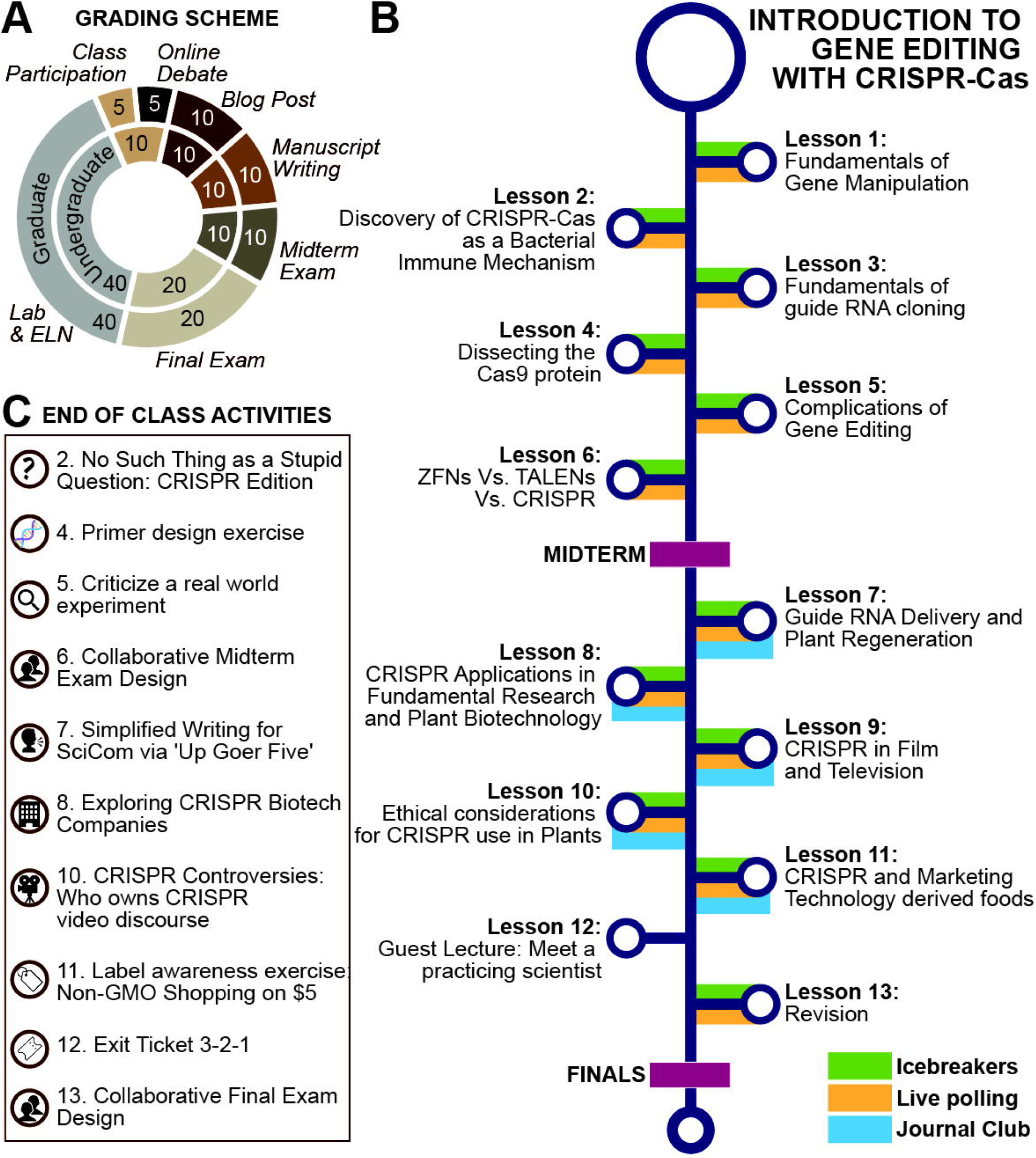
Lesson plan overview **A**, Grading scheme used in this course to avoid over reliance on any one form of testing throughout the semester. **B**, Lesson plan for a typical 14-15 week course taught during a university semester. Colored bars indicate the type pf activity undertaken during the class. C, List of end of class activities paired with relevant lessons in panel B.

**Figure 2.**
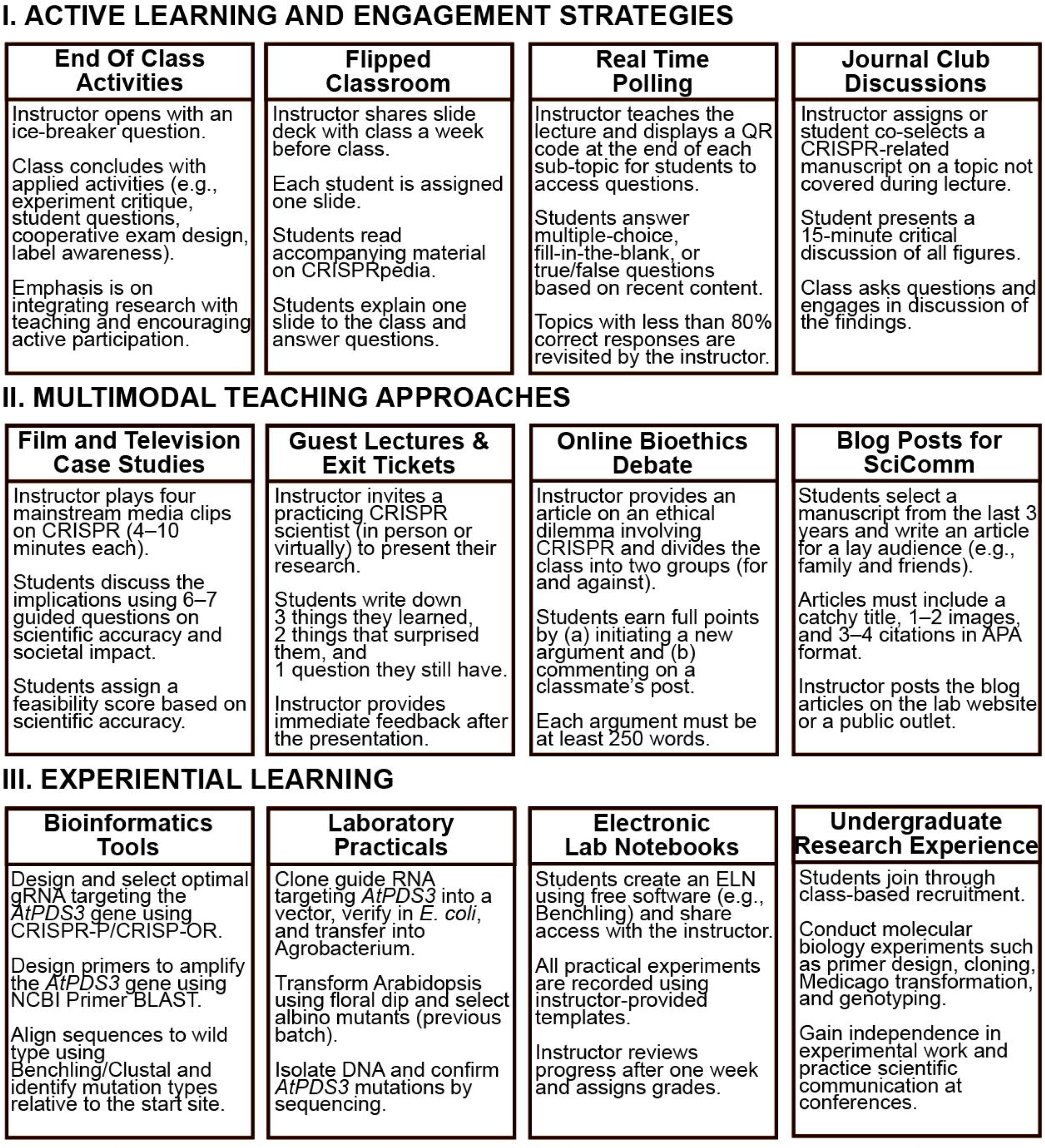
Overview and synopsis of teaching strategies used in this course to promote student engagement with subject material.

In addition to lectures, the course incorporated a variety of activities designed to enhance student engagement through active learning, multimodal instruction, and hands-on practicals (**Figure 1)**.

### 3.1 Active Learning and Engagement Strategies

Below, we outline the active learning and engagement classroom strategies employed: (1) End-of-class activities to reinforce concepts (2) Flipped classroom sessions (3) Real-time quizzes (4) Journal club discussions and homework assignments.

#### 3.1.1 End / start of class activities

End- or start-of-class activity sessions provided a rapid reinforcement for the knowledge gained and gave students the opportunity to apply those concepts in real time, boosting understanding and memory (Arthurs & Kreager, 2017). The following activities were employed.

1. Critiquing Real-World Experimental Designs. Students were encouraged to identify flaws, limitations, or potential improvements in published experiments or from the instructor’s own research, deepening their understanding of gene editing approaches and encouraging critical analysis of scientific literature.
2. No such thing as a stupid question. Guided by the instructor, students were encouraged to ask questions freely—no matter how simple or unusual they seemed. This approach helped make curiosity feel natural, removed any hesitation around asking questions, and gave instructors the chance to clear up confusion right away.
3. Cooperative Exam Design: Students formulated potential exam questions based on the lecture content to address what should another student who took this course know? They were encouraged to include conceptual and application-based questions. Selected questions were discussed and sometimes included in actual assessments. This activity reinforced content while promoting deeper engagement.
4. Label awareness: Students browsed online food and asked to add to their shopping carts foods that were labelled Non-GMO and comparable foods that were unlabelled. This led to a discussion about how labelling foods as Non-GMO might affect consumer choices.
5. Start-of-class ice breaker questions, such as “what superpowers would you like to get if you could edit your genome using CRISPR” helped build a warm collegial environment.
6. Students were asked to select only five plant species to take to a new planet and justify their choices from a set that included Arabidopsis, cotton, rice, soybean, oak, sunflower, sage, banana, potato, and Artemisia. This exercise prompted reflection on the diverse roles plants play in human life, ranging from food and medicine to cultural and religious value and challenged students to differentiate between essential and non-essential needs.

#### 3.1.2 Flipped classroom for selected lectures

In a “Flipped Classroom’, students engage with the material beforehand – by reading, watching videos, and building a foundation only to transform the classroom into a dynamic space for discussions, presentations, and collaborative learning with the professor as a guide.

A modified flipped classroom approach was implemented in the CRISPR course wherein each student was required to present one slide to their fellow classmates from a slide deck prepared by the instructor and shared a week beforehand. Each slide was assigned to a different student, making every participant responsible for researching and presenting a specific subtopic. Students delivered their material in 5-10-minute sessions, followed by a - minute period dedicated to instructor-guided questions and discussion. This structure encouraged all students to actively participate and engage with the content, as they not only prepared their own topics but also reviewed the entire subject to gain a broader context/understanding of CRISPR and its diverse applications. The use of CRISPR-pedia derived literature in this lecture (Lecture 4) provided students with a starting point for their research and helped them learn the basics of the technology. Compared to traditional lecture-based methods, this approach transformed students from passive listeners into active contributors to the learning process.

#### 3.1.3 Real time quizzes using polling software

To enhance engagement and evaluate learning in real time, the instructor used real-time quizzes in each lecture. If fewer than 70% of students answered a question correctly, the instructor revisited the material, and concepts were explained again before new topics were covered. This software provided a tool to assess both attendance and understanding among students throughout the semester.

#### 3.1.4. Journal club discussions and homework

Students selected or were assigned research articles to present to the class, including aspects of CRISPR-Cas technology that had not been covered in depth in class, applications, and foundational articles. Students had the flexibility to structure their own presentations, with the primary guideline being a focus on the interpretation of figures and panels from the selected manuscript. The article was shared in advance with the entire class, and all students were expected to read it before the session. At the beginning of each class, a 15–20-minute journal club presentation followed by a group discussion was held, allowing for peer-led engagement and critical analysis prior to the instructor’s lecture. The discussions primarily centered on the application of molecular tools to achieve precise gene editing. Interpreting graphs and figures in CRISPR-Cas-related publications, initially a challenge, became progressively clearer through the structured journal club sessions. These discussions enhanced comprehension of experimental design, methodologies, and data interpretation while keeping up to date with a fast-paced research field like CRISPR-Cas.

### 3.2 Multimodal teaching approaches

Multimodal teaching incorporates different modalities such as audio and visual inputs. To enhance the learning experience, this course incorporated: (1) Film and television clips (2) Guest lectures (3) Online ethics debate and (4) Blog posts.

#### 3.2.1 Film and television as fictional case studies

The integration of auditory, visual and interactivewnsiveness in STEM education (Tang, 2014; Hosťovecký & Štubňa, 2015; Öcal *et al*., 2021; Marozzo *et al*., 2024). Short film clips from *Rampage* (2018), *Coroner: Crispr Sistr* (2019), *Gattaca* (1997), *Spider-Man* (2002), and were used in this class to stimulate discussion about the applications of CRISPR gene editing technology. After watching the clips, students were asked to act as scientific consultants and score the scientific soundness of the clip based on a set of guided questions. These questions were related to (1) Feasibility of phenotype based on one or multiple gene acquisitions (2) CRISPR repair pathways (3) Nature of the target genetic change specifically, whether it resulted from an insertion or deletion, (4) Existence and feasibility of this CRISPR technology in real world, (5) Delivery method of the guide RNA-cas9 system, (6) Identifying the villain or ethical conflict -whether it is a scientist, science itself or another entity, (7) Portrayals of the potential social impacts of CRISPR, and (8) Feasibility score based on their overall assessment.

#### 3.2.2 Guest lectures and exit ticket exercises

Guest lectures by practicing scientists offer real-world insights and cover specific topics typically not included in regular classes, connecting students to the frontiers of the field, and demonstrating how new knowledge is generated. This approach improves the learning experience for students, teachers, and guest speakers (Ma, 2025). In this course, guest lectures covered CRISPR reagent delivery using nanoparticles, CRISPR applications in artificial intelligence, and the marketing challenges of GMOs and edited plants. These guest lectures introduced new teaching styles and expanded mentorship networks while increasing student attentiveness.

In addition to these teaching lectures, the course also invited two practicing plant biologists applying CRISPR in plant biology who presented their research projects where students performed an ‘Exit Ticket’ exercise. An exit ticket is a short writing assignment prompting students to answer questions based on lecture content. Like quizzes, exit tickets help teachers monitor students’ understanding and provide insights into their learning shortcomings, encouraging practice and reducing anxiety (Rodriguez *et al*.). During guest lectures, students were given an ‘exit ticket’ sheet at the beginning of the class containing three prompts: “Three things you learned,” “Two things that surprised you,” and “One question that still puzzles you” (**Supplementary Material 1**). This activity helped to focus students’ attention and provided an opportunity for the instructor to correct any misunderstandings in a timely manner.

#### 3.2.3 Online bioethics debates using eLearn

Students are often unprepared to engage in discussions about the real-world scientific and ethical dimensions of the material presented in their courses (Tibell & Rundgren, 2010; Seiter & Fuselier, 2021). Ethical considerations are often neglected in STEM classrooms, perhaps because they are viewed as peripheral to the “hard science” content or because they raise uncomfortable societal questions. However, it is essential that students develop the ability to critically evaluate how emerging biotechnologies should be used, taking into account their potential effects on human communities, ecological systems, and future generations. Today’s students will become the scientists responsible for advancing these tools, communicating their risks and benefits to the public, and shaping decisions that may influence millions of lives.

In the CRISPR course at TSU, students participated in an asynchronous online debate on the real-life case of Dr. He Jiankui, who controversially edited human embryos to resist HIV (Krimsky, 2019; Raposo, 2019) (**Supplementary Material 1**). While his stated intentions appeared benevolent, which was highlighted by students in favor of human gene editing, students identified critical ethical violations, including the absence of informed consent, secrecy, and disregard for international guidelines. The asynchronous format enabled participants to research, reflect, and compose thoughtful responses. Their arguments ranged from concerns over off-target effects and mosaicism to the potential misuse of science under social and political pressures.

#### 3.2.4 Blog posts for science communication

Blog posts are short, informal essays published on online platforms, where authors can express their perspectives, concepts, and feelings (Mansor, 2011; Wang & Chiou, 2022; Mbila-Uma *et al*., 2024). Since blogging spreads knowledge, feedback, communication, and collaboration, it is a strong and effective student-centered teaching tool for university-level science students (Blau *et al*., 2009; Wang & Chiou, 2022; Obionwu *et al*., 2023). Students in biological sciences or genetic engineering need conceptual and problem-solving education rather than mere knowledge of facts (Knight & Wood, 2005), and writing blog posts can provide this type of learning. Writing blog posts encourages students to review current and past research on relevant topics and provide their own critical thoughts on the subject of study.

In the TSU CRISPR course, students were tasked with writing a blog on the topic “**CRISPR Breakthroughs: Unveiling the Future of Genetic Engineering**” (**Supplementary Material 1**). The TSU College of Agriculture’s media and magazine editor was invited to discuss the key elements of a good blog post. The editor reviewed the basic characteristics of effective blog writing and provided structure and tips. Students were instructed to focus on a specific CRISPR breakthrough, analyzing and articulating its implications and potential future developments. They were encouraged to include at least two visual elements (images, infographics, diagrams) with captions and five credible sources, including at least two peer-reviewed scientific articles. The best blog posts were featured on the instructor’s lab website.

### 3.3 Experiential learning

In addition to the different approaches to teaching and learning described above, this course also included laboratory and computational exercises and research experiences. Specifically, students participated in: (1) Hands-on, research-based practicals to reinforce experimental design and techniques (2) Use of bioinformatics tools to design guide RNAs (gRNAs) and detect CRISPR-induced mutations (3) Electronic lab notebooks (ELNs) for documenting practical work, promoting organization and reproducibility (4) Undergraduate research experiences to expose students to real-world scientific inquiry.

#### 3.3.1 Hands-on lab practicals

The “Introduction to Gene Editing with CRISPR-Cas” course was designed as a hands-on educational experience, equipping students with both theoretical understanding and practical skills in gene editing. The hands-on practical lessons used CRISPR-Cas guideRNA cloning to demonstrate techniques such as polymerase chain reaction, gel electrophoresis, Type IIS restriction enzyme-based cloning, and the creation of targeted mutations that they could subsequently perform in their own research (Course Structure Overview – **Table 1**).

For this course, the phytoene desaturase (*PDS*) gene of *Arabidopsis thaliana* was selected as the target. Mutations in the *PDS* gene result in an albino phenotype (Qin *et al*., 2007), causing plants to appear white, thus providing a visually distinguishable marker for identifying mutants versus non-mutants (Mayta *et al*., 2022).

#### 3.3.2 Activities performed by the students

Briefly, students were introduced to bioinformatics tools that allowed them to obtain the gene of interest (*Arabidopsis thaliana PHYTOENE DESATURASE 3*) and design guide RNAs capable of introducing mutations into it, and PCR primers that clone and generate the guide RNA as well as the targeted gene to determine if the editing had been successful (detailed protocols of all steps are included in the **Supplementary Materials 2 and 3**).

In the wet-lab exercises, students used PCR to amplify and clone the guide RNA into bacterial cells, then sequenced the plasmids to ensure that the correct sequences had been cloned. The plasmids were then used to transform *Agrobacterium tumefaciens*, the microbial vector that introduces DNA into plant cells. Students transformed both wild-type *Arabidopsis thaliana* plants and mutants in *rdr* (RNA-dependent DNA Polymerase mutant plants are more susceptible to gene editing) via the floral dip method. Although the students did not characterize the transformants that they generated due to the long developmental timeframe of plants, they knew that they were generating samples for the subsequent year’s students to analyze.

In parallel, students were provided with seeds of previously generated wild-type and *pds3* mutant *A. thaliana* plants for genotyping. Students plated seeds from wild-type plants as well as plants that had been edited in the *pds3* gene. They grew the plants for four weeks, at which point they were able to visually identify wild-type (green), albino homozygous (white) or chimeric (green and white) plants. They selected control plants as well as edited plants with various phenotypes for DNA extraction. The students designed primers to amplify *AtPDS3*, characterized the product by gel electrophoresis, and sent the amplified PCR products out for sequencing. All students were able successfully genotype and analyze the edits in their mutant plants.

#### 3.3.3 Use of electronic lab notebooks to document practical work

Contemporary laboratories have shifted from handwritten notes to electronic data management systems, recognizing the benefits of digital record-keeping, including better organization, global accessibility, and improved efficiency in data retrieval and analysis (Kanza *et al*., 2017; Chen *et al*., 2024; Khan-Trottier, 2024). An Electronic Laboratory Notebook (ELN) digitizes traditional lab notebook record-keeping, allowing users to record protocols, observations, and experimental notes seamlessly. Researchers can access this data from computers and mobile devices, enhancing organization, documentation, collaboration, and data sharing https://ari.hms.harvard.edu/electronic-laboratory-notebooks-elns-guidelines

In the CRISPR course at Tennessee State University, the online electronic lab notebook Benchling was used. Benchling is a free platform for academic users that provides 10GB of storage. Importantly, it allows sharing with instructors for corrections and grading. Benchling includes molecular biology tools valuable for sequence analysis.

Students were required to document all practical aspects of their experiments in Benchling. The retrieved DNA sequences were uploaded to Benchling and the built-in tools used to design guide RNAs and primers. The instructor was able to check the students’ work and provide feedback as needed. The electronic notebook was an important part of the assessment strategy. As technology progresses, ELNs are expected to evolve, increasing their usability and positively impacting laboratory efficiency. Despite initial challenges in adoption and the assumption that all attendees must have a computer, ELNs have become an indispensable tool in advanced molecular biology teaching, research and documentation.

#### 3.3.4 Undergraduate research experiences

This course also served as a pipeline to recruit talented undergraduate students into the instructor’s laboratory on legume functional genetics. CRISPR-Cas-mediated genome editing offers unprecedented opportunities to rapidly develop crop varieties with improved yield, quality, and resilience to both biotic and abiotic stresses (Tuncel *et al*., 2025). Recognizing its transformative potential in advancing sustainable agriculture, undergraduate research projects were strategically designed to provide hands-on experience that not only reinforces classroom learning but also builds the innovative thinking and practical skills essential for success in future scientific careers. At TSU, undergraduate research students work on a flexible schedule and are compensated hourly in accordance with university-established wage rates. Those maintaining a GPA above 3.0 may also be eligible to affiliate their research with the Dean’s Scholars Program, a distinction that enhances their curriculum vitae and contributes to their academic and professional development. The projects built on the knowledge students acquired during classroom lectures and extended them further into fundamental research. For example, the projects offered over the two years of this course were (1) Testing effects of *MtBAK1 (BRASSINOSTEROID ASSOCIATED KINASE1)* multiplex editing on nodulation in *Medicago truncatula* (2) Genotyping transgenic Medicago plants edited in the peptide *MtCAPE16 (CAP-DERIVED PEPTIDE 16)*.

Some students carried out one-semester of research, but most stayed for two or more, and the projects were designed to fit their schedule. Additionally, students were encouraged to apply for funding to attend research conferences and present their work. Over the past two years, students presented their research at ARD Biennial Symposium in Nashville, TN, the TSU Research Symposium, the MANRRS 39th Annual National Conference in Memphis, TN, and the American Society of Plant Biology (ASPB) meeting in Milwaukee, WI, In addition to presenting their research posters, conference attendance helps students learn to engage effectively in networking sessions craft clear and concise self-introductions for professional and academic settings.

### 3.4 Student assessment techniques

This course used multiple and varied assessment metrics which avoided the over-reliance on exams as a measure of student learning (**Figure 1A**). Diverse assessment techniques provided fairer assessments of student learning; this course employed exams, class participation, participation in online activities, participation in practical activities and documentation in the electronic laboratory notebook, in addition to self-assessment/ self reflection on the part of the instructors to continuously improve the course based on feedback.

## 4 DISCUSSION

In comparison to traditional teaching approach, these interactive methods put greater emphasis on interaction, rapid feedback and student input in learning, all of which has shown to enhance learning outcomes for the STEM education system (Prince, 2004). Importantly, many of these strategies were implemented with minimal additional resources, using freely available digital tools or simple classroom modifications, making this model both sustainable and scalable.

These interactive activities gave some tangible and intangible outcomes. First, students developed a strong sense of critical thinking concrete learning and solid conceptual basis of fundamentals of gene editing as shown by the end of the course survey where 50% of students felt more confident in their understanding of CRISPR concepts as a result of the active learning sessions compared to 26% who self-reported no change and 20% who said their confidence reduced (**Figure 3A**). Intangibly these teaching methods helped build a stronger sense of teamwork in the classroom, improved the connection between students and the teacher, and encouraged students to believe they could grow and improve. These factors are key to helping students succeed in challenging science subjects.

**Figure 3.**
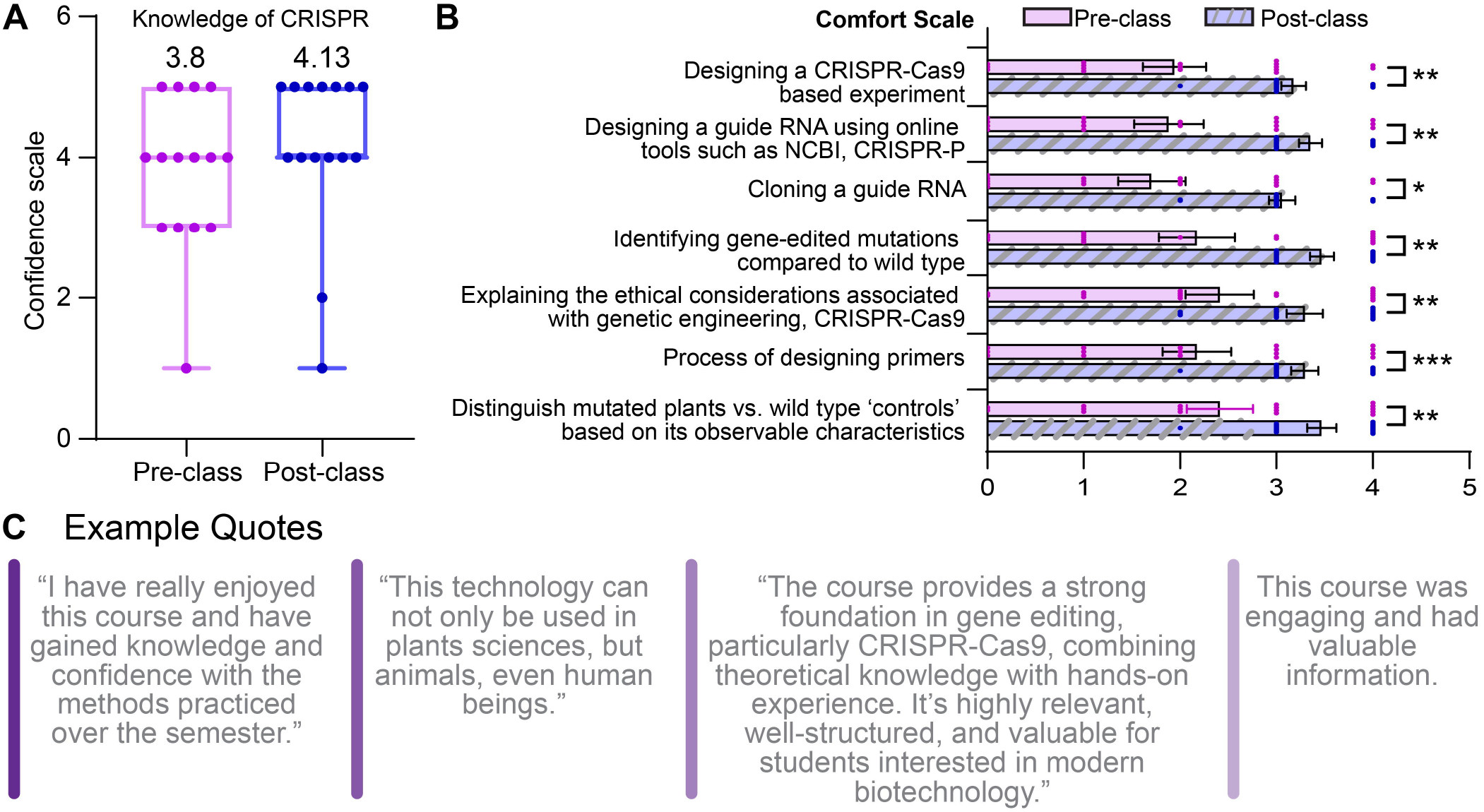
Student survey responses across two semesters. **A.** Pre- and post-course matched survey responses to the statement “I feel confident about my knowledge of CRISPR-Cas9 in plant science/agriculture.” Data represent 15 of 21 students who completed both surveys. **B.** Pre- and post-course matched survey responses measuring student comfort with the hands-on components of the course. Each data point indicates the comfort score and error bars represent SEM. Statistical significance was determined using Student’s t-test (*p*<0.05, ^*^*p*< 0.01,^* *^ *p* < 0.001). n = 17 students who completed both surveys.

The key benefit that these approaches offer is in their ability to promote active and student-centered learning. Unlike general lecture setup where students are just passive receivers of information these activities promote knowledge construction, peer collaboration, and real-time application of concepts. These teaching methods help in tackling the problem of students feeling bored or disconnected in STEM associated classes. By adding fun, low-pressure activities during each of the learning sessions, students got regular chances to dive into the material in a way that worked for them.

The flipped classroom model made the topic of CRISPR more approachable and engaging for students. By fostering active participation and collaboration, this approach enhances interaction between students and teachers, creating a supportive learning environment where students feel confident to ask questions and focus on understanding complex concepts. Furthermore, this teaching method significantly contributes to curriculum improvement by motivating students to advance their research efforts, explore topics more deeply, and cultivate curiosity and critical thinking.

While the “Introduction to Gene Editing with CRISPR-Cas” course was highly effective in a small classroom setting, scaling it for larger cohorts presents several logistical challenges. High levels of instructor-student interaction, a cornerstone of the engagement strategy, are time-intensive and may be difficult to sustain with increased enrolment. Lab sessions occasionally extended beyond the allotted time due to limited access to specialized equipment, such as clean benches required for sterile work. Additionally, the course was dependent on a teaching assistant for preparing lab materials—an institutional support not universally available. The use of time-sensitive reagents also required precise scheduling and testing, which may not be feasible in all instructional settings. Therefore, careful adaptation of course modules based on available resources will be paramount in implementing such a course at different institutions.

Though developed within the context of plant sciences, this curriculum, which is grounded in molecular biology, can be adapted for broader fields such as animal or food sciences. Lessons specifically focused on plant transformation and agricultural applications (e.g., Lessons 7 and 8) can be modified to suit other biological systems, and modules such as bioethics can be refocused on human or animal contexts. Commercial kits from Bio-Rad and the Innovative Genomics Institute offer accessible alternatives for demonstrating CRISPR-Cas9 activity, but they can be cost-prohibitive for some institutions. In contrast, the use of *Arabidopsis* as a teaching organism offers a cost-effective and renewable platform for hands-on learning, especially when reagents like SapI, ligase, and primers are purchased in bulk and properly stored. Competent cells can be prepared in-house and preserved at−80°C, further minimizing recurring costs.

Ultimately, this course demonstrates that exposing students to cutting-edge STEM technologies like CRISPR, when combined with active learning and hands-on experiences significantly enhances their engagement, builds confidence, and cultivates practical skills. Given that this manuscript was written by students who took this course can arguably be seen as evidence of engagement as well. Such course designs are essential for preparing a future-ready, scientifically literate workforce equipped to contribute meaningfully to molecular and biotechnological research.

## ACKNOWLEDGEMENTS

We thank the guest speakers for their valuable contributions and expertise shared with the class. We would also like to thank Steve Jacobson and Zheng Li of UCLA for sharing their *pds3* mutant lines with us. We thank Elizabeth Haswell, The Sustainable Professor for careful reading and feedback on the manuscript.

## AUTHOR CONTRIBUTION

This work was supported by NSF awards #2205542 and #2217830 and USDA-NIFA grant 2022-38821-37353, awarded to Sonali Roy. S.R. conceptualized the project and taught the course. D.J. and S.G.D. served as teaching assistants and contributed to the development of the experimental templates provided in the supplementary materials. R.T. collected anonymous student feedback and evaluations. V.C. assisted in designing assignments and assessment rubrics. M.W. reviewed and edited the manuscript and helped formulate exams. All student authors contributed equally to the manuscript text and are listed in alphabetical order. All authors wrote, read and approved the final version of the manuscript.

## DATA AVAILABILITY STATEMENT

Data sharing not applicable to this article as no datasets were generated or analyzed during the current study. Materials described in the manuscript are included as Supplementary materials and can be used according to journal policies or a CC BY-NC-SA license.

## REFERNCES

Alaagib NA, Musa OA, Saeed AM. 2019. Comparison of the effectiveness of lectures based on problems and traditional lectures in physiology teaching in Sudan. BMC Medical Education 19(1): 365.

Ansori AN, Antonius Y, Susilo RJ, Hayaza S, Kharisma VD, Parikesit AA, Zainul R, Jakhmola V, Saklani T, Rebezov M, et al. 2023. Application of CRISPR-Cas9 genome editing technology in various fields: A review. Narra J 3(2): e184.

Arthurs LA, Kreager BZ. 2017. An integrative review of in-class activities that enable active learning in college science classroom settings. International Journal of Science Education 39(15): 2073–2091.

Blau I, Mor N, Neuthal T. 2009. Open the Windows of Communication: Promoting Interpersonal and Group Interactions Using Blogs in Higher Education. Interdisciplinary Journal of E-Learning and Learning Objects 5.

Chen SH, Garcia CB, Lazear E, Lentz TB, Robertson SD, Li Z, Goller CC. 2024. Implementation of electronic lab notebooks (ELNs) in science laboratory classes: student response and lessons learned. Discover Education 3(1): 70.

Freeman S, Eddy SL, McDonough M, Smith MK, Okoroafor N, Jordt H, Wenderoth MP. 2014. Active learning increases student performance in science, engineering, and mathematics. Proceedings of the National Academy of Sciences 111(23): 8410–8415.

Hosťovecký M, Štubňa J 2015. The Impact of Film-based Learning in Science Education.In Elleithy K, Sobh T. New Trends in Networking, Computing, E-learning, Systems Sciences, and Engineering. Cham: Springer International Publishing. 531–536.

Hubbard K. 2024. Plant biology education: A competency-based vision for the future. PLANTS, PEOPLE, PLANET 6(4): 780–790.

Jinek M, Chylinski K, Fonfara I, Hauer M, Doudna JA, Charpentier E. 2012. A Programmable Dual-RNA–Guided DNA Endonuclease in Adaptive Bacterial Immunity. Science 337(6096): 816–821.

Kanza S, Willoughby C, Gibbins N, Whitby R, Frey JG, Erjavec J, Zupančič K, Hren M, Kovač K. 2017. Electronic lab notebooks: can they replace paper? Journal of Cheminformatics 9(1): 31.

Khan-Trottier A. 2024. Use of OneNote class notebook as a combined electronic laboratory notebook and content delivery tool in an introductory biochemistry laboratory course. Biochemistry and Molecular Biology Education 52(4): 462–473.

Knight JK, Wood WB. 2005. Teaching more by lecturing less. Cell Biol Educ 4(4): 298–310.

Krimsky S. 2019. Ten ways in which He Jiankui violated ethics. Nature Biotechnology 37(1): 19–20.

Ma M. 2025. Enhancing Student Engagement and Learning Outcomes Through Strategic Use of Guest Speakers in Advertising Education. Journal of Advertising Education 29(1): 41–52.

Mansor AZ. 2011. Reflective Learning Journal Using Blog. Procedia - Social and Behavioral Sciences 18: 507–516.

Marozzo V, Crupi A, Abbate T, Cesaroni F, Corvello V. 2024. The impact of watching science fiction on the creativity of individuals: The role of STEM background. Technovation 132: 102994.

Mayta ML, Dotto M, Orellano EG, Krapp AR. 2022. An experimental protocol for teaching CRISPR/Cas9 in a post-graduate plant laboratory course: An analysis of mutant-edited plants without sequencing. Biochemistry and Molecular Biology Education 50(5): 537–546.

Mbila-Uma S, Koleoso O, Umoga I, Alassad M, Agarwal N. 2024. The Multi-attribute impact of hyperlinks in blogs: an emotion-centric approach. Social Network Analysis and Mining 14(1): 134.

Mills A, Jaganatha V, Cortez A, Guzman M, Burnette JM, Collin M, Lopez-Lopez B, Wessler SR, Norman JMV, Nelson DC, et al. 2021. A Course-Based Undergraduate Research Experience in CRISPR-Cas9 Experimental Design to Support Reverse Genetic Studies in Arabidopsis thaliana. Journal of Microbiology & Biology Education 22(2): 10.1128/jmbe.00155-00121.

Obionwu CV, Broneske D, Saake G 2023. Microblogs-A means for simulating informal learning beyond classrooms. Proceedings of the 14th International Conference on Education Technology and Computers. Barcelona, Spain: Association for Computing Machinery. 219–225.

Öcal En, Yildirim B, Sahin Topalcengiz E. 2021. STEM IN MOVIES: FEMALE PRESERVICE TEACHERS’ PERSPECTIVES ON MOVIE “HIDDEN FIGURES”. Journal of Baltic Science Education 20: 740–758.

Olson S, Riordan DG. 2012. Engage to excel: producing one million additional college graduates with degrees in science, technology, engineering, and mathematics. Report to the president. Executive office of the president.

Prince M. 2004. Does Active Learning Work? A Review of the Research. Journal of Engineering Education 93(3): 223–231.

Qin G, Gu H, Ma L, Peng Y, Deng XW, Chen Z, Qu L-J. 2007. Disruption of phytoene desaturase gene results in albino and dwarf phenotypes in Arabidopsis by impairing chlorophyll, carotenoid, and gibberellin biosynthesis. Cell Research 17(5): 471–482.

Raposo VL. 2019. The First Chinese Edited Babies: A Leap of Faith in Science. JBRA Assist Reprod 23(3): 197–199.

Rodriguez M, le Roux C, Melville M. Iteratively-Designed Exit Tickets Enhances Student Learning. College Teaching: 1–9.

Seiter KM, Fuselier L. 2021. Content knowledge and social factors influence student moral reasoning about CRISPR/Cas9 in humans. Journal of Research in Science Teaching 58(6): 790–821.

Sudarmika P, Santyasa I, Divayana DG. 2020. Comparison between Group Discussion Flipped Classroom and Lecture on Student Achievement and Student Characters. International Journal of Instruction 13: 171–186.

Tang M. 2014. The impact of science fiction media on student interest and learning. Montana State University.

Tibell LAE, Rundgren C-J. 2010. Educational Challenges of Molecular Life Science: Characteristics and Implications for Education and Research. CBE—Life Sciences Education 9(1): 25–33.

Tuncel A, Pan C, Clem JS, Liu D, Qi Y. 2025. CRISPR–Cas applications in agriculture and plant research. Nature Reviews Molecular Cell Biology: 1–23.

Wang Y-M, Chiou C-C. 2022. Empirically Examining the Effectiveness of Teaching Blogs on University Course Instruction. Sage Open 12(3): 21582440221104782.

Wolyniak MJ, Austin S, Bloodworth LF, Carter D, Harrison SH, Hoage T, Hollis-Brown L, Jefferson F, Krufka A, Safadi-Chamberlin F, et al. 2019. Integrating CRISPR-Cas9 Technology into Undergraduate Courses: Perspectives from a National Science Foundation (NSF) Workshop for Undergraduate Faculty, June 2018. Journal of Microbiology & Biology Education 20(1): 10.1128/jmbe.v1120i1121.1702.

Yuan P, Usman M, Liu W, Adhikari A, Zhang C, Njiti V, Xia Y. 2024. Advancements in Plant Gene Editing Technology: From Construct Design to Enhanced Transformation Efficiency. Biotechnology Journal 19(12): e202400457.

Zeng H, Shu X, Wang Y, Wang Y, Zhang L, Pong TC, Qu H. 2021. EmotionCues: Emotion-Oriented Visual Summarization of Classroom Videos. IEEE Transactions on Visualization and Computer Graphics 27(7): 3168–3181.

